# Haspin kinase binds to a nucleosomal DNA supergroove

**DOI:** 10.1101/2024.05.21.595243

**Authors:** Chad W. Hicks, Colin R. Gliech, Xiangbin Zhang, Sanim Rahman, Stacy Vasquez, Andrew J. Holland, Cynthia Wolberger

## Abstract

Phosphorylation of histone H3 threonine 3 (H3T3) by Haspin recruits the chromosomal passenger complex to the inner centromere and ensures proper cell cycle progression through mitosis. The mechanism by which Haspin binds to nucleosomes to phosphorylate H3T3 is not known. We report here cryo-EM structures of the Haspin kinase domain bound to a nucleosome. In contrast with previous structures of histone-modifying enzymes, Haspin solely contacts the nucleosomal DNA, inserting into a supergroove formed by apposing major grooves of two DNA gyres. This unique binding mode provides a plausible mechanism by which Haspin can bind to nucleosomes in a condensed chromatin environment to phosphorylate H3T3. We identify key basic residues in the Haspin kinase domain that are essential for phosphorylation of nucleosomal histone H3 and binding to mitotic chromatin. Our structure is the first of a kinase domain bound to a nucleosome and is the first example of a histone-modifying enzyme that binds to nucleosomes solely through DNA contacts.

## Introduction

Post-translational modification of histones regulates fundamental biological processes, including transcription, DNA replication, the DNA damage response, and the cell cycle ^1^. Histone phosphorylation regulates the cell cycle by triggering a cascade of histone modifications to promote chromatin condensation during mitosis ^2,3^. Phosphorylation of histone H3 threonine 3 (H3T3ph) and histone H2A threonine 120 (H2AT120ph) define the inner centromere and are two of the most prominent mitotic phosphoryl modifications ^4^. H3T3ph, which is established at the nuclear envelope in early prophase, becomes most concentrated at inner centromeres during metaphase, and gradually diminishes during anaphase ^5^. The Repo-Man/PP1γ complex dephosphorylates H3T3ph to oppose the spread of this modification to chromosome arms in early mitosis and catalyzes the removal of H3T3ph at the end of mitosis ^6^. H3T3ph recruits the chromosomal passenger complex (CPC) to inner centromeres ^4,7,8^. The CPC, one of the main controllers of mitosis ^9^, contains the highly conserved Aurora B kinase, which phosphorylates multiple histone H3 residues ^10^ and many kinetochore proteins ^11^ at the centromere.

The Haspin kinase establishes and maintains wild-type levels of H3T3ph during mitosis^12,13^ and is conserved in most eukaryotes, including yeast, plants, flies, and worms ^14,15^. The only known substrate of Haspin is histone H3T3. Depletion or inhibition of Haspin abolishes H3T3ph ^13^, displaces the CPC from centromeres ^16^, causes premature chromatid separation ^17^, prevents normal chromosome alignment ^13^, and delays mitotic progression ^16,18^. The BUB1 kinase can partially compensate for loss of Haspin by phosphorylating histone H2A-T120, which helps recruit the CPC to inner centromeres ^19^ ^20,21^. Although earlier studies have argued that the H3T3 kinase, VRK1, is also important for generating H3T3ph during mitosis ^22,23^, more recent work suggests that VRK1 is not essential for generating mitotic H3T3ph, while Haspin is essential ^24^.

Human Haspin is a 798 amino acid protein that contains a ∼450 residue disordered N-terminal region and a C-terminal atypical kinase domain ^25–28^. Haspin’s structured C-terminal kinase domain has a bilobal shape similar to that of proteins belonging to the eukaryotic protein kinase (ePK) family ^14,27,28^, one of the largest families of eukaryotic proteins ^29^. Haspin contains a negatively charged active-site cleft between its two structured lobes where the histone H3 tail and adenosine triphosphate (ATP), the phosphate donor, bind ^28,30^. Mono-, di-, and tri-methylation of histone H3 lysine 4 (H3K4me1/2/3), which is commonly found at actively transcribed genes ^31^, inhibit Haspin activity ^28^, likely though steric clash of the methylated lysine within the enzyme’s acidic active site cleft ^30^. Since Haspin is important for ensuring proper mitotic progression ^12,13^ and is overexpressed in many cancers ^32^, there have been extensive efforts to identify and characterize compounds that inhibit Haspin H3T3 phosphorylation activity ^28,33–36^.

Whereas the structural basis by which Haspin recognizes the histone H3 tail is well-characterized ^27,28,30^, nothing is known about how this enzyme engages an intact nucleosome substrate. While many chromatin-binding proteins contact the nucleosome acidic patch and other residues in the histone octamer core ^37–39^, these surfaces are often occluded in a condensed chromatin environment. It is not known how Haspin binds to nucleosomes in these conditions.

We report here the structure of the Haspin kinase domain bound to a nucleosome determined by cryogenic electron microscopy (cryo-EM). Unlike all previous structural studies of nucleosome binding proteins, Haspin does not contact the globular histone core. Instead, Haspin exclusively contacts the DNA, inserting into a cavity formed by apposed major grooves of two adjacent gyres of nucleosomal DNA. Haspin engages the unique geometry of this DNA “supergroove” ^40^ with basic residues that contact the negatively charged DNA sugar-phosphate backbone. The positioning of Haspin in the supergroove between superhelical locations (SHL) 5.5 and -2.5 enables Haspin to bind the histone H3 tail and engage its H3T3 substrate. Our structure, the first of any histone kinase domain bound to chromatin, reveals a novel mode of nucleosome binding that can allow Haspin to phosphorylate its histone substrate in both open and condensed chromatin.

## Results

### Cryo-EM structure of Haspin bound to a nucleosome

To investigate the structural basis by which Haspin binds nucleosomes, we used cryo-EM to determine structures of the Haspin kinase domain (residues 465-798) bound to nucleosome core particles wrapped with 185 bp of DNA containing the 147 bp Widom 601 sequence flanked by 19 bp linkers (185bp) **(Table 1, Figure S1)**. Samples were prepared by mixing Haspin with nucleosome in the absence of crosslinker and flash-freezing on cryo-EM grids **(see Methods)**. The cryo-EM density maps revealed Haspin bound to the side of the nucleosome disk, engaging both gyres of nucleosomal DNA **(Figure 1A-B)**. We identified two similar but distinct orientations of Haspin, denoted positions 1 and 2 **(Figure 1C).** The global resolution estimates for the EM maps of Haspin in positions 1 and 2 were 3.01 Å and 2.99 Å, respectively **(Figures S2A-B)**. The local resolution of Haspin in position 1 is ∼4 Å and ∼5 Å in position 2 **(Figures S2C-D)**. Within the EM density for each Haspin kinase domain, the resolution is highest at the interface between Haspin and the nucleosome. The fit for the Haspin model in position 1 was slightly better than for Haspin in position 2, but the secondary structure of both Haspin molecules fits well within their respective EM maps **(Figures S2E-F)**. The structure of the Haspin kinase domain is very similar to previously published crystal structures **(Figure S3)** ^27,28,30^, indicating that there are no significant conformational changes in Haspin upon binding to nucleosome.

**Figure 1.**
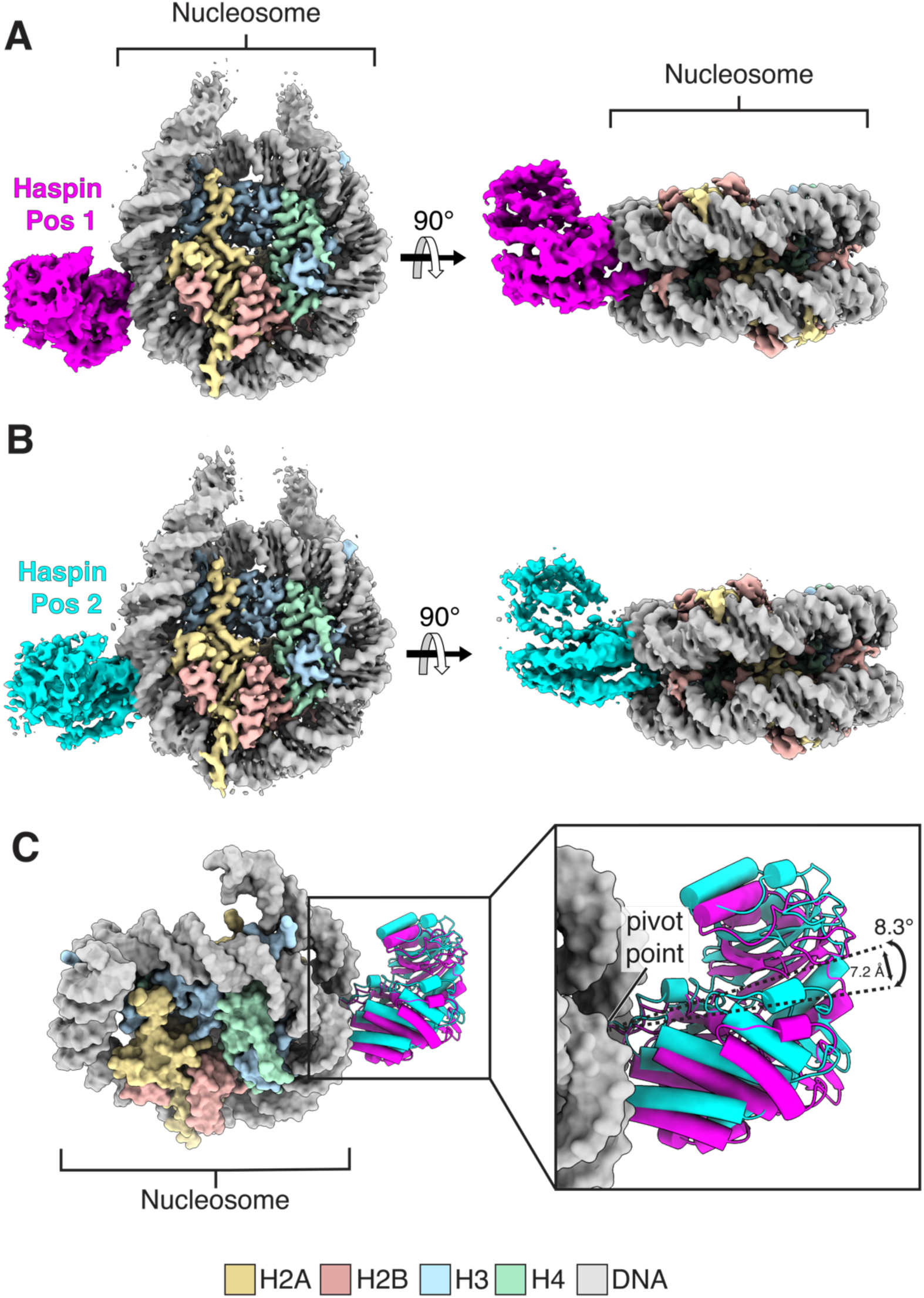
Haspin binds to nucleosomal DNA. **A)** Cryo-EM map of Haspin (465-798) in position 1 bound to nucleosome **B)** Cryo-EM map of Haspin (465-798) in position 2 bound to nucleosome. **C)** Superposition of the nucleosomes for cryo-EM models of Haspin bound to nucleosome in position 1 and 2. Haspin cartoon models show an 8.3° pivot between the two Haspin positions with the pivot point origin located at the SHL 5.5 major groove.

**Table 1.** Cryo-EM details and validation metrics. Table of details related to the two cryo-EM structures of Haspin bound to nucleosome with Haspin in position 1 and 2, and a third cryo-EM structure of locally-refined Haspin. The details include parameters for data collection, data processing, model refinement, and model validation.

|  | Haspin<br>position 1 | Haspin<br>position 2 | Haspin local<br>refinement |
| --- | --- | --- | --- |
| **Data Collection and Processing** |  |  |  |
| Magnification (X) | 130,000 | 130,000 | 130,000 |
| Voltage (kV) | 300 | 300 | 300 |
| Electron exposure (e<sup>-</sup>/Å<sup>2</sup>) | 40 | 40 | 40 |
| Dose rate (e<sup>-</sup>/px/s) | 7.74 | 7.74 | 7.74 |
| Defocus range (μM) | -0.5 to -2.5 | -0.5 to -2.5 | -0.5 to -2.5 |
| Pixel size (Å) | 0.970 | 0.970 | 0.970 |
| Camera | Falcon 4 | Falcon 4 | Falcon 4 |
| Energy Filter slit width (eV) | 10 | 10 | 10 |
| Micrographs (#) | 9,362 | 9,362 | 9,362 |
| Initial cleaned particle stack (#) | 900,411 | 900,411 | 900,411 |
| Particle duplication symmetry | --- | --- | --- |
| Final particles (#) | 152,199 | 154,151 | 481,127 |
| Map resolution at 0.143 FSC cutoff (Å) | 3.01 | 2.99 | 3.64 |
| **Refinement** |  |  |  |
| Initial models used (PDB) | Nuc: 4ZUX<br>Haspin: 4OUC | Nuc: 4ZUX<br>Haspin: 4OUC | H3: 4OUC<br>Haspin: 4OUC |
| Map resolution range (Å) | 2.00 to 6.00 | 2.00 to 6.00 | 2.00 to 10.00 |
| Model Resolution at 0.5 FSC cutoff (Å) | 3.2 | 3.2 | 4.3 |
| Non-hydrogen atoms (#) | 15,497 | 15,194 | 2,672 |
| Nucleotides (#) | 314 | 314 | 0 |
| Ligands (#) | 0 | 0 | 0 |
| Protein B factor (Å<sup>2</sup>) | 84.34 | 116.92 | 135.69 |
| Nucleotide B factor (Å<sup>2</sup>) | 102.40 | 131.09 | --- |
| Ligand B factor (Å<sup>2</sup>) | --- | --- | --- |
| Bond length RMSD (Å) | 0.005 | 0.004 | 0.004 |
| Bond angles RMSD (°) | 0.618 | 0.583 | 1.081 |
| **Validation** |  |  |  |
| MolProbity score | 1.67 | 1.80 | 1.97 |
| Clashscore | 7.27 | 7.94 | 14.78 |
| Rotamer Outliers (%) | 0.52 | 2.24 | 0.67 |
| Ramachandran Favored (%) | 96.08 | 97.52 | 95.74 |
| Ramachandran Allowed (%) | 3.92 | 2.50 | 4.26 |
| Ramachandran Disallowed (%) | 0.00 | 0.00 | 0.00 |

### Haspin binds to a nucleosome supergroove

Haspin binds to nucleosomes via a unique DNA-binding mechanism. Unlike all other structures of histone-modifying enzymes bound to nucleosome reported to date ^37–39^, the Haspin kinase domain does not contact the histone core. Instead, the enzyme binds to nucleosomal DNA by inserting into a large cavity formed by the apposition of major grooves on the two neighboring gyres of nucleosomal DNA, termed a “supergroove” ^40^ **(Figure 2A)**. There are multiple such supergrooves at different locations around the nucleosome core particle. The specific supergroove to which Haspin binds is composed of the major grooves at superhelical locations (SHL) 5.5 and -2.5. **(Figure 2A)**. To our knowledge, this is the first example of any protein that binds to a nucleosome in this manner.

**Figure 2.**
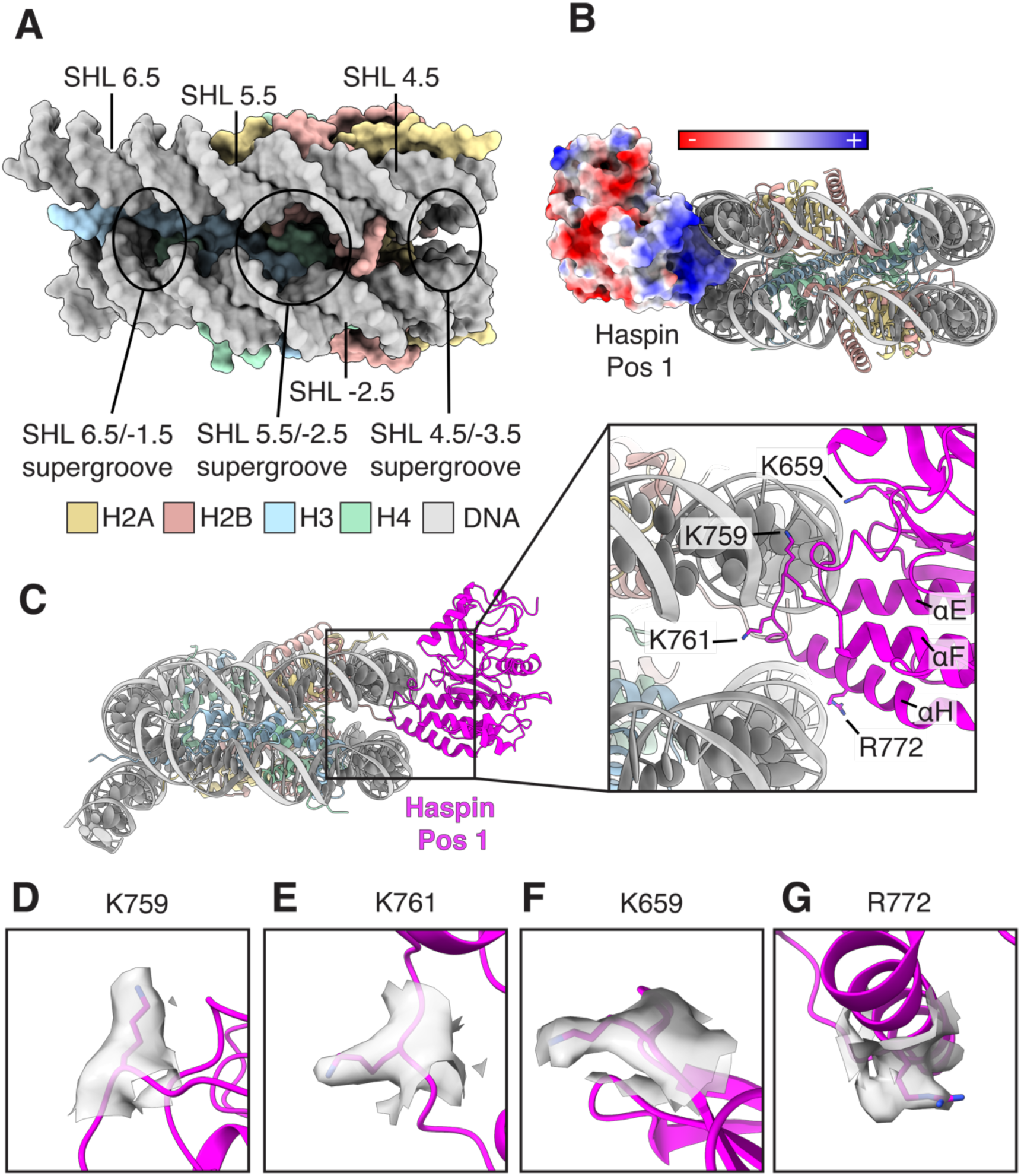
Extensive ionic interactions stabilize Haspin binding to the nucleosomal DNA supergroove. **A)** Nucleosome surface representation showing how DNA major grooves on adjacent nucleosomal DNA gyres form nucleosomal DNA pores at different super-helical locations (SHLs). **B)** Surface model of Haspin bound to nucleosome in position 1 colored in electrostatic representation. **C)** Cartoon model of Haspin bound to nucleosome in position 1 showing key Haspin residues interacting with nucleosomal DNA. **D-G)** Cryo-EM side-chain density for key Haspin residues (K759, K761, K659, R772).

Extensive electrostatic interactions between the positively charged surface of the Haspin kinase domain and the negatively charged DNA stabilize Haspin binding to the nucleosome **(Figure 2B)**. The positive ends of the Haspin αE and αH helix dipoles insert into the DNA supergroove **(Figure 2C)**, while residues K759 and K761 in the loop connecting helices αF and αH form electrostatic interactions with the sugar-phosphate backbone on either side of the major groove at SHL 5.5 **(Figure 2C-E)**. In addition, K659 forms electrostatic interactions with the sugar-phosphate backbone adjacent to SHL 5.5 **(Figure 2C,F),** and R772 interacts with the DNA phosphate backbone adjacent to SHL -2.5 **(Figure 2C,G)**. Side chain and main chain atoms of K759 and T760 are sufficiently close to DNA bases at positions 55 and -56 to form van der Waals interactions. We note that the DNA sequence in this location is not identical in the twofold symmetry-related nucleosomal DNA, but the density is ambiguous, suggesting that our maps likely represent an average of particles containing Haspin bound to one or the other symmetry-related supergrooves. There are otherwise no apparent contacts with the DNA bases.

The identification of two distinct Haspin positions on the nucleosome suggests that there is some flexibility in the positioning of Haspin within the supergroove. Superposition of the nucleosomes in the two models corresponding to Haspin positions 1 and 2 shows the kinase domain rotates by ∼8.3° about a pivot point located in the DNA major groove at SHL 5.5 **(Figure 1C)**. The direction of the rotational change in Haspin position follows the curve of the major groove at SHL 5.5. Although Haspin contacts both DNA gyres, the rotational change in enzyme position about a pivot point at SHL 5.5 suggests that interactions with the SHL 5.5 DNA gyre play a more dominant role in positioning Haspin than interactions with the SHL -2.5 DNA gyre.

The end of the H3 tail in the Haspin active site was poorly resolved, likely due to heterogeneity in the position of Haspin bound to nucleosome **(Figure 1C)**. To better resolve the H3 tail, we removed the EM density corresponding to nucleosome from the cryo-EM particle stack using signal subtraction and performed local refinement and alignment of the Haspin kinase domain alone **(Table 1, Figure S1)** to a final resolution of 3.64 Å **(Figure S4A)**. The local resolution of the Haspin local refinement ranged from ∼2 Å at the center of the EM map to ∼10 Å at the periphery **(Figures S4B)**, allowing a better Haspin model fit at the center of EM map than at the periphery **(Figure S4C)**. We observed well-resolved EM density for H3 residues 2-5 showing a 180° turn of the H3 tail, positioning H3R2, H3T3, and H3K4 deep within the Haspin acidic cleft **(Figure 3A-B)**.This is consistent with a previously determined crystal structure of Haspin bound to H3 tail peptide (1-7) showing a similar binding position and turn in the H3 tail (PDB: 4OUC) ^30^ **(Figure S5A-B).**

**Figure 3.**
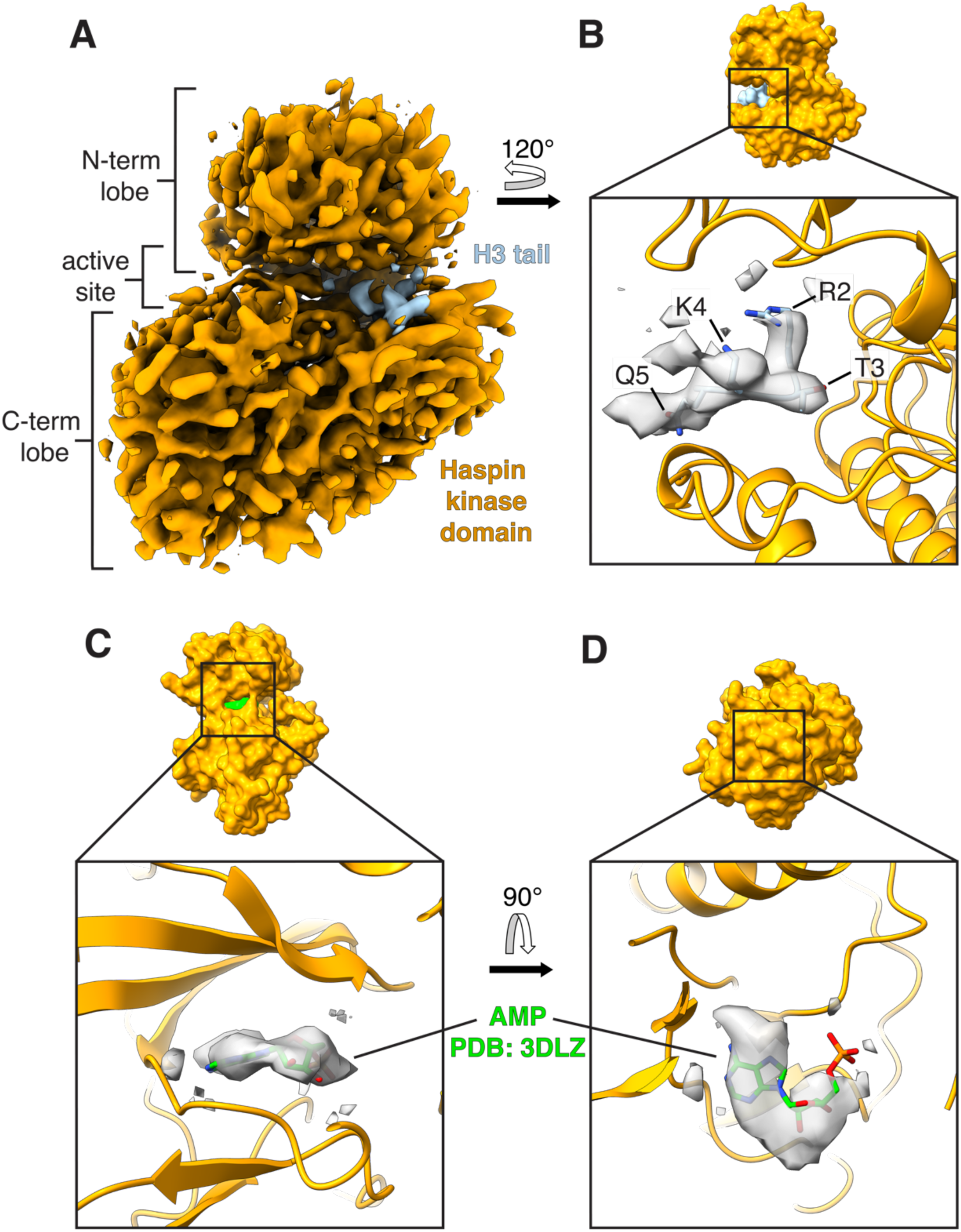
Local Refinement of Haspin reveals the H3 tail and a small molecule bound to the active site. **A)** Cryo-EM map of locally-refined Haspin (465-798) from a cryo-EM dataset of Haspin bound nucleosome. **B)** Cryo-EM surface and cartoon models of Haspin (465-798) showing H3 tail bound to the active site with good EM density for H3 tail side-chains (2-5). **C-D)** Cryo-EM surface and cartoon models of Haspin (465-798) showing density for a small molecule bound to the Haspin ATP cofactor site. A previously reported Haspin structure (PDB: 3DLZ) with bound AMP molecule was superimposed over our cryo-EM map of Haspin and fits well within our EM density.

Although no ATP or nucleotide analogs were added during sample preparation, we also observed density for a small molecule in the Haspin ATP binding site **(Figure 3C-D)**. We presume that the density corresponds to a small molecule that bound to Haspin during its overexpression and remained bound during purification. Superposition of the crystal structure of Haspin kinase domain bound to adenosine monophosphate (AMP) (PDB: 3DLZ) ^28^ shows a good fit of an AMP molecule to the EM density **(Figure 3C-D).**

While we did not observe strong EM density for the histone H3 tail in our initial unsharpened cryo-EM maps **(Figure 1A-B)**, low-pass filtering the map of Haspin bound to nucleosome in position 1 revealed low-resolution density for the H3 tail **(Figure 4A)**. The H3 tail follows a path along the nucleosomal DNA, bridging the gap to Haspin’s structured N-terminal kinase domain lobe, and inserting into the Haspin active site **(Figure 4B)**. While individual residues cannot be resolved in this map, this portion of the H3 tail contains multiple positively charged residues that could form favorable electrostatic interactions with the negatively charged DNA backbone **(Figure 4C)**. Whereas our model shows one discrete representative model for the path of the H3 tail along the nucleosome, the low resolution of the EM density for the tail suggests that these residues are highly mobile and can adopt multiple conformations along its path to the Haspin active site.

**Figure 4.**
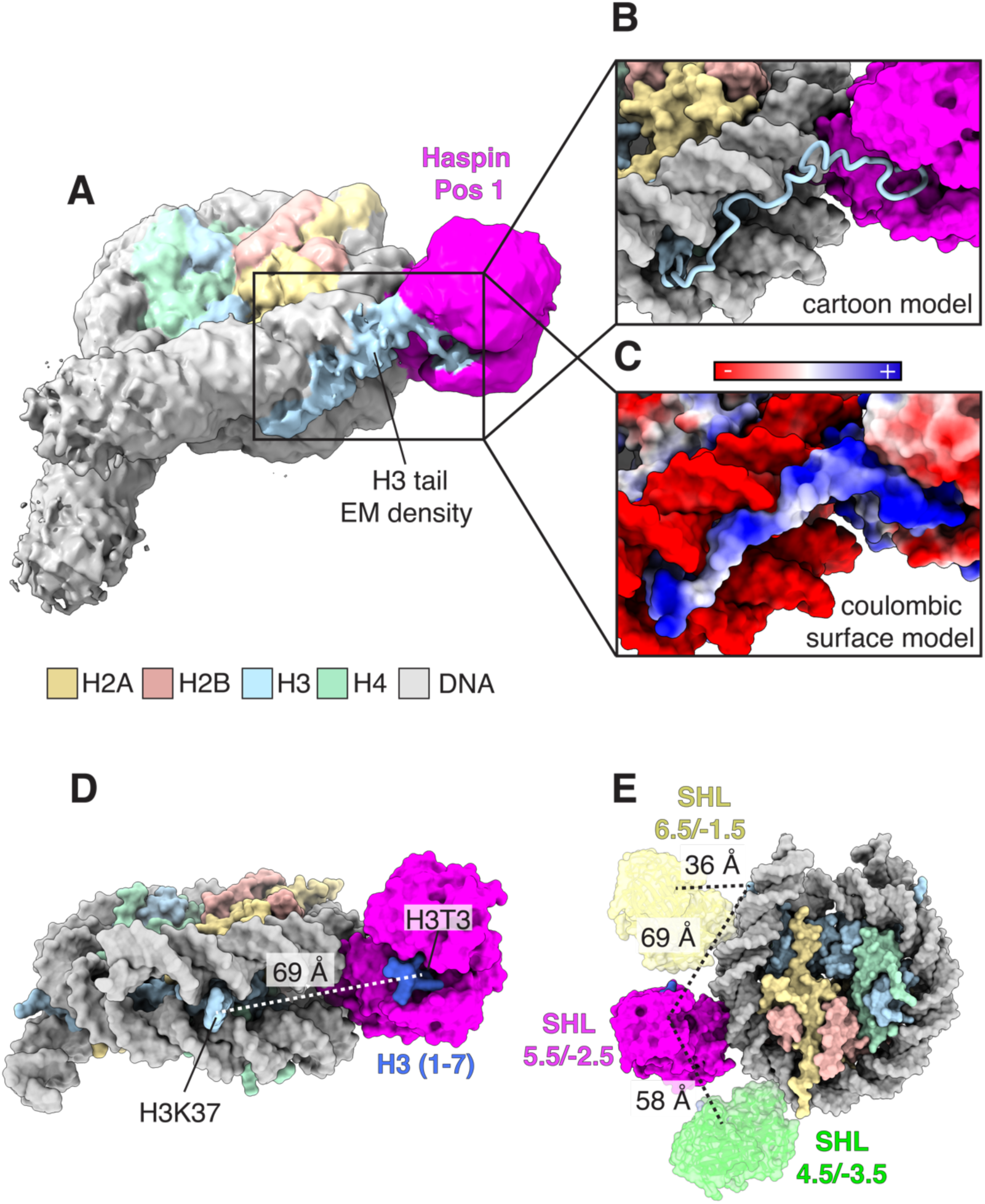
Haspin binds in an optimal position for engaging the H3 tail. **A)** Gaussian filtered cryo-EM map of Haspin bound to nucleosome in position 1 at a very low threshold showing EM density for the H3 tail. **B)** Atomic model of Haspin bound to nucleosome in position 1 depicting a path for the histone H3 tail into the Haspin active site. **C)** Cryo-EM surface model colored in coulombic surface representation. **D)** Superimposition of the cryo-EM model of Haspin in position 1 and the crystal structure of Haspin bound to H3 tail peptide (dark blue) (PDB: 4OUC)(Maiolica et al., 2014) showing a 69 Å distance from the H3T3 and the last well-ordered H3 tail residue in the cryo-EM model (residue H3K37) (light blue). **E)** Distance calculations between H3K37 of the cryo-EM structure of Haspin bound to nucleosome and H3T3 of hypothetical Haspin binding positions in the SHL 6.5/-1.5 or SHL 4.5/-3.5 supergroove.

Binding to the SHL 5.5/-2.5 DNA supergroove would place the Haspin active site ∼69 Å from H3K37, the last well-ordered histone H3 residue, which emerges between the DNA gyres **(Figure 4D)**. Since the maximum distance that can be covered by H3 residues 3-37 is ∼119 Å, binding to the SHL 5.5/-2.5 DNA supergroove positions Haspin at an optimal distance to bind the extended histone H3 tail with Thr3 in the enzyme active site. Binding to the adjacent major supergroove at SHL 4.5/-3.5 would place the Haspin active site ∼127 Å from H3K37, **(Figure 4E)**, which would position Haspin too far away to bind to H3T3. Although binding to the supergroove at SHL 6.5/-1.5 would place the Haspin active site only ∼36 Å from H3K37, well within reach of H3T3 **(Figure 4E)**, we did not observe EM density for Haspin in this supergroove. We note that the H3 tail would need to form a long loop or fold upon itself to place H3T3 within the enzyme active site at the SHL 6.5/-1.5 supergroove. A less extended conformation of the H3 tail would likely reduce the potential to form ionic interactions with the DNA, making this a less-favored binding site for Haspin.

Since the 2.5/-5.5 supergroove is symmetrically analogous to the 5.5/-2.5 supergroove, Haspin should be able to bind to both supergrooves, which would result in particles corresponding to 2:1 Haspin:nucleosome complexes. However, our analysis of the cryo-EM data revealed particles with just one Haspin kinase domain bound to a nucleosome, or no bound enzyme at all. We speculate that the affinity of Haspin for nucleosomes is not high enough to have produced a significant number of 2:1 Haspin:nucleosome complexes at the Haspin and nucleosome concentrations used for cryo-EM sample preparation. Consistent with this explanation, 47% of nucleosomes in the cryo-EM data were not bound by Haspin **(Figure S1)** and needed to be removed using 3D classification to obtain a stack of particles containing only nucleosomes with bound Haspin.

### Basic residues are important for Haspin binding and activity *in vitro* and in cells

To evaluate the contributions of individual Haspin residues to nucleosome binding, we assayed the effects of point substitutions on the affinity of Haspin for nucleosomes using electrophoretic mobility shift assays (EMSA). Increasing concentrations of wild-type Haspin resulted in the formation of a discrete shifted band, which presumably corresponds to a 1:1 complex of Haspin bound to nucleosome **(Figure 5A)**. At the highest concentrations of Haspin, there appeared a high molecular weight smear resulting from non-specific aggregates of Haspin and nucleosome. In binding assays of Haspin point mutants containing the charge-reversal substitutions, K659E, K759E, K761E, or R772E, the discrete band corresponding to a complex of Haspin bound to nucleosome is lost **(Figure 5A)**. Introducing all four substitutions in Haspin abrogated all detectable binding to nucleosomes, even at an enzyme concentration of 32 μM **(Figure 5A)**.

**Figure 5.**
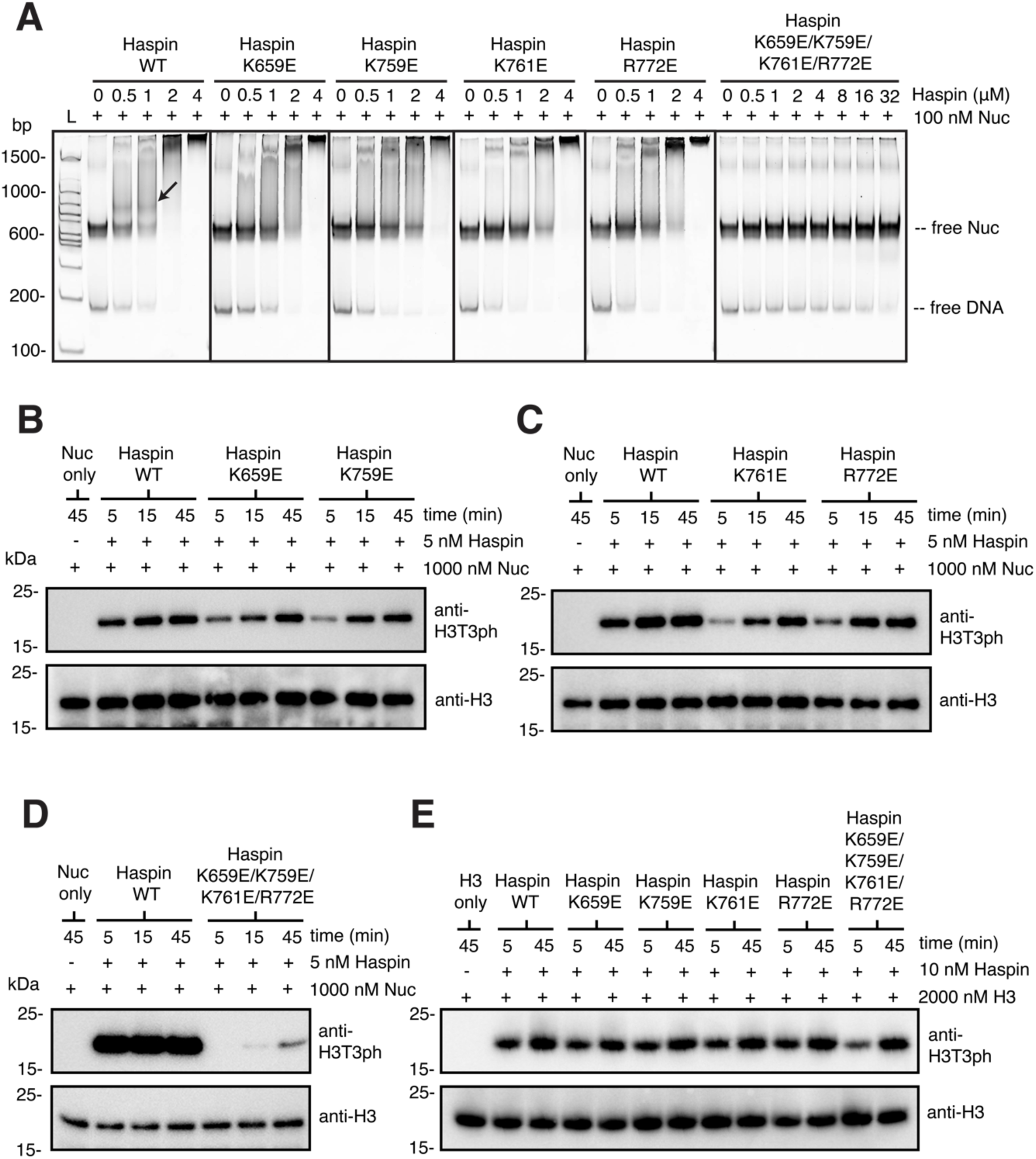
Charge-reversal mutagenesis of key Haspin residues disrupts Haspin binding to nucleosome and H3T3 phosphorylation activity. **A)** Electrophoretic mobility shift assays (EMSAs) showing binding of wild-type Haspin (465-798) and Haspin mutants (465-798) to nucleosome at the indicated concentrations. Black arrow points to band indicative of a Haspin-nucleosome complex. **B-D)** Immunoblots showing activity of wild-type Haspin and Haspin mutants on nucleosomes. **E)** Immunoblot showing activity of wild-type Haspin and Haspin mutants on free histone H3. Activity was measured by incubating with an antibody against H3T3ph. Loading controls were run on a separate gel and incubated with an antibody against histone H3.

To confirm the relevance of the observed Haspin-DNA contacts to kinase activity, we assayed the effects of Haspin mutations on phosphorylation of nucleosomal histone H3T3. Point substitutions K659E, K759E, K761E, and R772E in Haspin all reduced the rate of H3T3 phosphorylation as compared to wild-type enzyme **(Figure 5B-C)**, Incorporating all four charge-reversal substitutions into Haspin dramatically reduced activity **(Figure 5D)**. These substitutions had no effect on the ability of Haspin to phosphorylate free histone H3 **(Figure 5E)** confirming that the mutations affect nucleosome binding only, not enzymatic activity.

We next tested whether the residues identified as important for DNA binding *in vitro* are also important for chromatin binding of full-length Haspin in cells. HEK293T cells were transfected with enhanced green fluorescent protein (EGFP)-tagged Haspin to assess its association with mitotic chromosomes. Consistent with previous reports, wild-type EGFP-Haspin co-localized with DAPI-stained DNA in fixed cell imaging **(Figure 6A)**. Strikingly, each of the four charge-reversal mutations (K659E, K759E, K761E, and R772E) caused a greater than four-fold reduction in the co-localization of Haspin with chromatin as compared to wild-type Haspin and was comparable to the EGFP-only control **(Figure 6B)**. Interestingly, some cells transfected with the Haspin mutant formed protein aggregates **(Figure 6A)**, which were not observed in cells transfected with wild-type Haspin. Taken together, these results support the importance of the DNA contacts observed in the cryo-EM structure of the Haspin kinase domain bound to a nucleosome.

**Figure 6.**
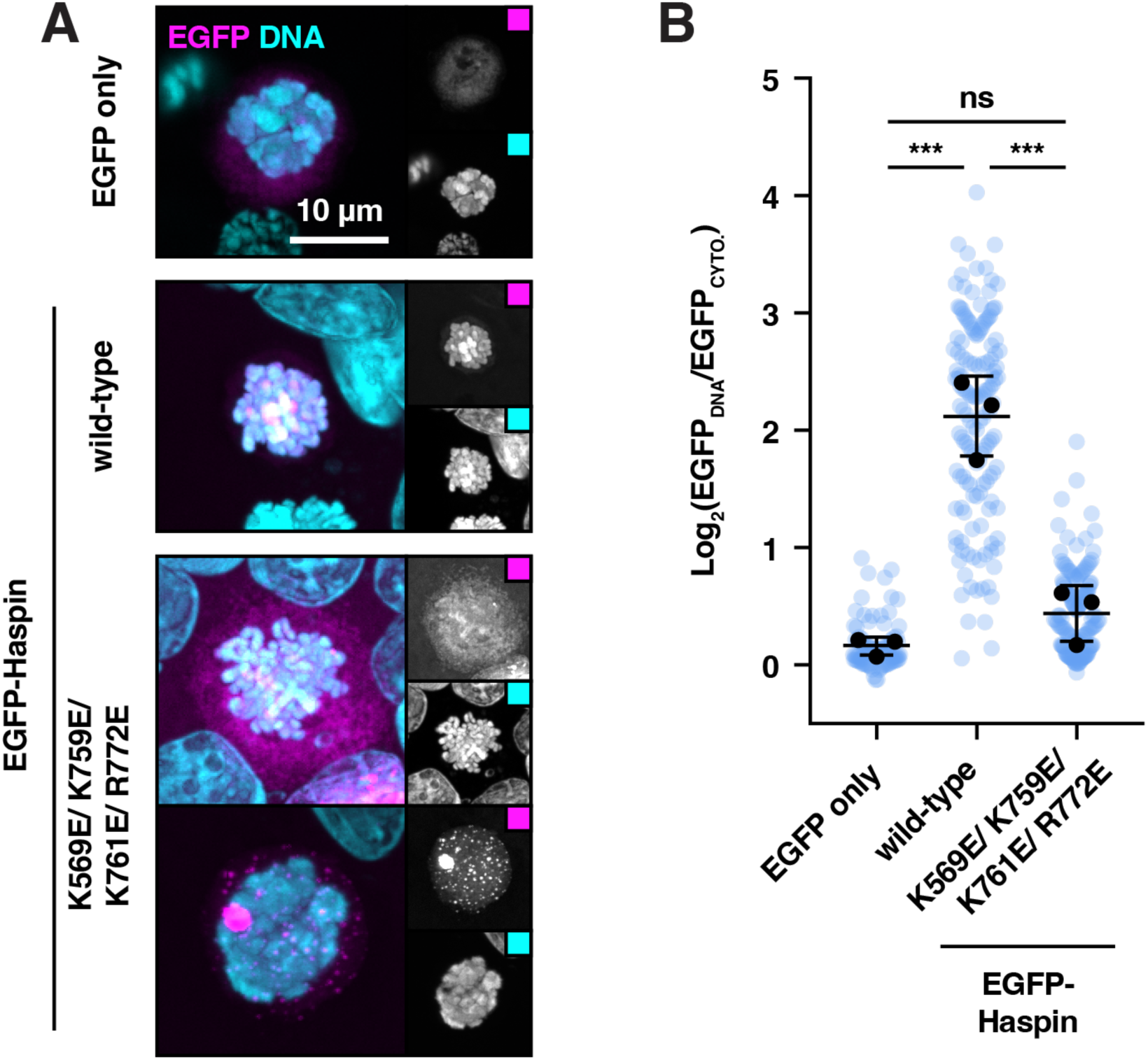
Positively-charged Haspin kinase domain residues are critically important for recruiting full-length Haspin to chromatin in cells. **A)** Fixed-cell fluorescence microscopy of mitotic HEK293T cells transfected with EGFP alone, EGFP-Haspin (wild-type), and EGFP-Haspin (K659E, K759E, K761E, and R772E). **B)** Ratio of cytoplasmic and DNA co-localized median EGFP signal from mitotic images.

## Discussion

Previous investigations had revealed the mechanism by which Haspin engages an H3 tail peptide ^30^, but it was not known how Haspin recognizes an intact nucleosome substrate. Our structure unexpectedly shows that the Haspin kinase domain binds solely to the nucleosomal DNA, inserting into the cavity called a supergroove ^40^ that is formed by two adjacent DNA major grooves **(Figures 1 and 2)**. To our knowledge, this is the first report of a protein that binds to this unique feature of nucleosomes and the first reported structure of a histone-modifying enzyme that does not contact the globular histone octamer core.

The potential of DNA supergrooves for specific interactions with proteins was first proposed by Edayathumangalam, Luger, and colleagues ^40^ in their study of a polyamide that binds to two adjacent minor grooves in the nucleosome, it was noted that the nucleosome contains multiple major and minor supergrooves, each arising from the apposition of DNA major and minor grooves on adjacent DNA gyres. While these supergrooves were proposed as potential sites that could be recognized by proteins that form sequence-specific base pair contacts within the supergroove ^40^, Haspin primarily contacts the sugar-phosphate backbone of the DNA and does not appear to recognize specific bases. This lack of specific interactions is consistent with the biological role of Haspin, which must bind to nucleosomes with diverse sequences.

The observed electrostatic interactions between the Haspin kinase domain and nucleosomal DNA are essential for Haspin binding to nucleosomes *in vitro* **(Figure 5)** and to chromatin in cells **(Figure 6)**. Previous studies had shown that Haspin can bind DNA ^25^ and speculated that the many positively charged residues in the unstructured N-terminus of Haspin mediated DNA binding ^14,25,26^. However, we have shown that charge-reversal substitutions of four key basic residues in the Haspin kinase domain almost completely abolish Haspin localization to mitotic chromatin in HEK293T cells **(Figure 6)**. While we cannot rule out the possibility that basic residues in the disordered N-terminal region of Haspin also contribute to chromatin binding, our results suggest that the observed interactions of the Haspin kinase domain with nucleosomal DNA is primarily responsible for the binding of this enzyme to chromatin.

Are there other proteins that bind to nucleosome supergrooves? A Foldseek^41^ search of the Alphafold database for structures similar to the C-terminal kinase domain lobe of Haspin (610-798) identified seven structurally related human proteins, all of which are kinases: AKT1, CLK2, MAPK1, MAPK14, CLK4, MAPK3, and PINK1. Whereas Haspin’s αH helix and αF-αH loop form extensive electrostatic interactions within the nucleosomal DNA supergroove (**Figure 2C**), all seven related proteins have a shorter αH helix and repositioned αF-αH loop that would not permit a mode of DNA binding similar to Haspin **(Figure 7A)**. In addition, none of the structural homologues has an electropositive surface that would be required to bind to negatively charged DNA **(Figure 7B)**. The ability of Haspin to bind to a major DNA supergroove thus appears to be an adaptation of the conserved kinase domain to bind nucleosomes. Future structural and bioinformatics investigations should help determine whether there are other chromatin-binding proteins that recognize the unique features of major and minor DNA supergrooves.

**Figure 7.**
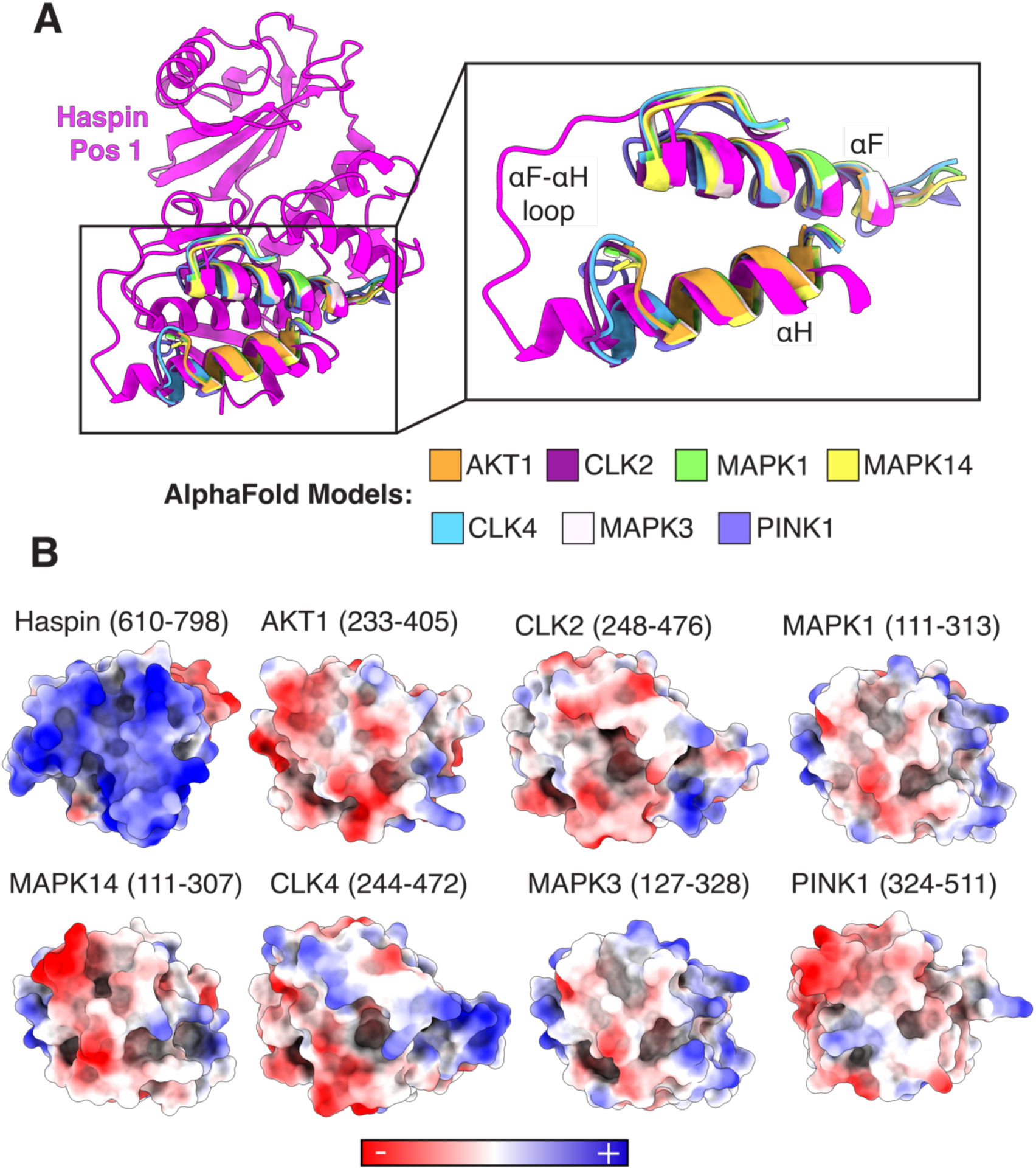
Haspin DNA supergroove binding surface surface is unique among structurally related proteins. **A)** Foldseek structural homology search using Haspin (610-798) as the search query and superimposition over Haspin. Inset shows Haspin (738-783), AKT1 (327-347,372-383), CLK2 (358-380,443-459), MAPK1 (205-227,281-294), MAPK14 (200-222,276-289), CLK4 (354-376,439-456), MAPK3 (223-244,298-311), PINK1 (430-452,476-490) in cartoon model representation. **B)** Superimposition of Haspin and structurally related proteins in coulombic surface representation showing the difference in charge of the Haspin DNA binding surface.

## Methods

### Expression and purification of histones

Expression plasmids for *Xenopus laevis* histones H2A, H2B, H3 and H4 were a gift from Greg Bowman (Johns Hopkins University). Histones were expressed and purified as previously described ^42^, with the following modifications. Thawed cell pellet was lysed with multiple passes through a Microfluidizer (Microfluidics). After lysis, centrifugation, and multiple washes using a Triton X-100 containing buffer, washed cell pellets were resuspended in a buffer containing 20 mM HEPES pH 7.5, 7 M Guanidine HCl, 10 mM dithiothreitol (DTT). Resuspended washed cell pellets were size exclusion purified using a buffer containing 10 mM Tris pH 8.0, 7 M urea, 1 mM EDTA, 5 mM b-mercaptoethanol (BME). Fractions were pooled and injected onto a 2-column series with a HiTrap Q-XL column connected upstream of a HiTrap SP-XL column using a buffer containing 20 mM Tris pH 7.8, 7 M urea, 1 mM EDTA, 5 mM BME). The Q-XL column was removed from the system, and then the histones were eluted from the SP-XL column with a gradient of 0 – 1 M NaCl on an ÄKTA Pure (Cytiva) chromatography system.

### Purification of Widom 601 DNA

The Widom 601 DNA sequence (147 bp) ^43^ was expressed in I strain XL-1 Blue using the pST55-16x601 plasmid ^44^. The 601 sequence was expressed, purified, and isolated as described ^45^. The 601 sequence used is:

5’−ATCGGATGTATATATCTGACACGTGCCTGGAGACTAGGGAGTAATCCCCTTGGCGGTTAAAACGCGGGGGACAGCGCGTACGTGCGTTTAAGCGGTGCTAGAGCTGTCTACGACCAATTGAGCGGCCTCGGCACCGGGATTCTCGAT−3’.

The Widom 601 DNA flanked by 19 bp of linker DNA at each end (185 bp) was amplified using polymerase chain reaction (PCR) using primers (IDT DNA) containing 19 bp overhanging regions using the linear double-stranded 147 bp 601 DNA as a template. One primer contained a biotin tag to allow for the amplification of biotinylated 185 bp DNA. The primers used were:

Forward:

5’−biosg−GTCGCTGTTCGCGACCGGCAATCGATGTATATATCTGACACGTGCC−3’

Reverse:

5’−GACCCTATACGCGGCCGCCCATCAGAATCCCGGTGCCGAG−3’

Phusion polymerase was used to amplify the reaction in 100 μL reaction volumes using standard PCR parameters. The PCR product mixture was precipitated by combining 100% EtOH, PCR product mixture, and 3 M sodium acetate pH 5.2 in a 10:1:20 ratio, and placing the mixture at -80°C for 1 hour. The precipitated PCR product mixture was then centrifuged and the supernatant removed. The pellet was washed with 70% EtOH, centrifuged, the supernatant removed, and the pellet allowed to dry. The pellet was resuspended in TE Buffer (10 mM Tris pH 8.0, 1 mM EDTA). A phenol:chloroform:isoamyl alcohol solution 25:24:1 was added to an equal volume of resuspended PCR product pellet, vortexed, and centrifuged to separate the phases. The organic phase was removed and the aqueous phase was extracted twice more the same way. All of the removed organic phases were pooled, back-extracted with TE buffer to collect any PCR product left behind in the organic phase, and all the aqueous phases combined and saved as the purified PCR product. Purified PCR product, 3 M sodium acetate pH 5.2, and 100% EtOH was combined in a 10 : 1 : 20 ratio and placed at -20°C overnight. The precipitated purified PCR product was then centrifuged, the pellet washed with 70% EtOH, centrifuged again, and allowed to dry. This pellet of purified 185bp 601 DNA was then resuspended in MilliQ (Sigma) water and stored at - 20°C.

### Preparation of nucleosomes

Unmodified nucleosome (185bp) was reconstituted as previously described ^42^, with the following modifications.

For each nucleosome sample, histone octamer and Widom 601 DNA was combined in an octamer:DNA molar ratio of 1.2 : 1 in a buffer containing 10 mM Tris pH 7.5, 2 M KCl, 1 mM EDTA, 1 mM DTT, such that the final DNA concentration was 6 μM. The salt concentration was gradually reduced to 0.25 M KCl over 24 hours by salt gradient dialysis to assemble nucleosome. Precipitate was removed by centrifugation and the purity of the nucleosome sample was assessed using an electrophoretic mobility shift assay (EMSA). If the nucleosome sample showed excess free DNA or higher-order species, it was further purified by loading onto a SK DEAE-5PW column (TOSOH biosciences) using a buffer containing 10 mM Tris pH 7.5, 0.25 M KCl, 0.5 mM EDTA, 1 mM DTT, and eluted with a gradient of 0.25 – 0.6 M KCl on an Agilent HPLC instrument. Purified nucleosome was dialyzed into a buffer containing 20 mM HEPES pH 7.5, 25 mM KCl, 1 mM EDTA, 1 mM DTT, 20% glycerol, flash frozen in liquid nitrogen, and stored at -80°C.

### Purification of Haspin and Haspin mutants

Haspin plasmid construct (GSG2) was a gift from Nicola Burgess-Brown (Addgene plasmid # 38915; http://n2t.net/addgene:38915; RRID:Addgene_38915). This construct was used to create Haspin mutant constructs using around-the-horn mutagenesis. The original Addgene wild-type Haspin construct or Haspin mutant constructs encoding Haspin (465-798) fused to an N-terminal hexahistidine tag (6xHis) and Tobacco Etch Virus (TEV) protease cut site, were transformed into BL21(DE3)Rosetta2-pLysS *E. coli* cells. The colonies were used to inoculate 5 mL volumes of media, which were expanded to larger 1 L volumes of media for the full-scale growth. Cultures were grown at 37°C and 200 RPM in 2x Yeast Extract Tryptone (2XYT) media supplemented with kanamycin and chloramphenicol. Cultures were induced by addition of 1 mM isopropyl-ß-D-thiogalactopyranoside (IPTG) when the culture reached an OD_600_ of 0.4 and were grown for an additional 16 hours at 18°C.The cells were harvested by centrifugation and the pellets resuspended in a buffer containing 20 mM HEPES pH 7.5, 300 mM NaCl, 40 mM imidazole, 5 mM BME, 1 tablet/ 50mL of complete protease inhibitor (Roche #11836153001), flash-frozen in liquid nitrogen, and stored at -80°C.

The frozen cell pellet suspensions were thawed in a water bath and an equal volume of buffer containing 20 mM HEPES pH 7.5, 300 mM NaCl, 40 mM imidazole, 1 mM DTT, and 0.2 mM PMSF was added. The diluted cell suspensions were lysed by sonication for three rounds each of 1 min total processing time (5 sec on/ 10 sec off) at 40% power. The resulting whole cell extracts were centrifuged at 17,000 RPM and the supernatants were filtered using a 1.1 μm filter to obtain clarified extracts. Clarified extracts were loaded onto a 5 mL HisTrap HP (Cytiva) column equilibrated in 20 mM HEPES pH 7.5, 300 mM NaCl, 40 mM imidazole, 1 mM DTT, and 0.2 mM PMSF and eluted with a gradient of 0.04 – 1 M imidazole on an ÄKTA pure instrument (Cytiva). The eluted proteins were diluted with 20 mM HEPES pH 7.5, 1 mM DTT, 0.2 mM PMSF to a final salt concentration of 150 mM NaCl and filtered using a 1.1 μm filter. Eluted Haspin mutant containing four charge-reversal mutations (K659E, K759E, K761E, and R772E) was diluted to a final salt concentration of 50 mM NaCl and filtered using a 1.1 μm filter. The filtered proteins were loaded onto a 5 mL HiTrap heparin (Cytiva) column equilibrated in 20 mM HEPES pH 7.5, 50 mM NaCl, 1 mM DTT, 0.2 mM PMSF and eluted with a gradient of 0.15 – 2 M NaCl on an ÄKTA pure instrument. The Haspin mutant containing for charge-reversal mutations was eluted from the heparin column with a gradient of 0.05 – 2 M NaCl. The eluted proteins were dialyzed overnight in a buffer containing 20 mM HEPES pH 7.5, 150 mM NaCl, 10% glycerol, 1 mM DTT, 0.2 mM PMSF, concentrated, flash-frozen in liquid nitrogen, and stored at -80°C.

### Cryo-EM sample preparation

Wild-type Haspin (465-798) containing an N-terminal 6xHis tag and TEV protease cut-site was thawed on ice and mixed with nucleosome containing 147 bp 601 Widom DNA with 19 bp linkers (185 bp) to a final concentration of 8 μM Haspin and 4 μM nucleosome in a buffer containing 20 mM HEPES pH 7.5, 150 mM KCl, 1 mM DTT, The mixture was incubated on ice for 30 min. Quantifoil R 1.2/1.3 gold 300 mesh grids (Electron Microscopy Sciences #Q350AR1.3) were glow-discharged for 45 seconds at 15mA using a PELCO easiGLOW Glow Discharge System to apply a negative charge to their surface. Then, 3 μL of sample mixture was applied to the grid, immediately blotted for 4.5 seconds with a blot force of 3, and plunge-frozen in liquid ethane using a Vitrobot Mark IV apparatus (Thermo Fisher) set to 100% humidity and 4°C.

### Cryo-EM data collection

To evaluate the feasibility of determining the cryo-EM structure of Haspin bound to nucleosome without using crosslinker, a small cryo-EM dataset of Haspin bound to nucleosome was collected at the Beckman Center for Cryo-EM at the Johns Hopkins University School of Medicine using a Thermo Fisher Glacios 200 kV electron microscope equipped with a Falcon 4i direct electron detector. A dataset of 1,026 exposures was collected in counting mode and recorded in Electron Event

Representation (EER) format using a magnification of 120kx, pixel size of 1.19 Å, nominal dose of 40 e^-^/ Å^2^, dose rate of 6.69 e^-^/px/s, and a defocus range of -0.5 to -2.5 μm. The final processed EM density map derived from this dataset was used to generate 2D templates for use during data processing of the publication-quality cryo-EM dataset of Haspin bound to nucleosome, which was collected using a Thermo Fisher Titan Krios 300 kV electron microscope.

Publication-quality cryo-EM data of Haspin bound to nucleosome was collected at the Beckman Center for Cryo-EM at the Johns Hopkins University School of Medicine using a Titan Krios at 300 kV equipped with a Falcon 4 Direct Electron Detector and Selectris Energy Filter. A dataset of 9,999 exposures was collected in counting mode and recorded in Electron Event Representation (EER) format using a magnification of 130kx, pixel size of 0.97 Å, nominal dose of 40 e^-^/ Å^2^, dose rate of 7.74 e^-^/px/s, a defocus range of -0.5 to -2.5 μm, and an energy filter slit width of 10 eV. A multi-shot imaging strategy was used to collect 3 shots per hole, utilizing beam image shift to move between each target.

### Cryo-EM data processing

The test dataset of Haspin bound to nucleosome collected on a Glacios 200 kV electron microscope was processed in cryoSPARC v4.2 ^46^. Exposures were imported with an EER upsampling factor of 2 and cropped to one-half of their original resolution using Patch Motion Correction. The contrast transfer function (CTF) correction was performed Patch CTF Correction. Poor quality micrographs were removed using Manually Curate Exposures, yielding 972 high-quality micrographs. An initial round of particle picking was performed using Blob Picker and Inspect Picks, then extracted using Extract from Micrographs. A set of 2D templates was created by first performing 2D classification on the extracted particles then selecting representative views using Select 2D classes. These templates were used to perform a second round of particle picking using Template Picker and Inspect Picks, then extracted using Extract from Micrographs to yield an uncleaned particle stack of 560,971 particles. One round of 2D Classification and Select 2D Classes was performed to remove junk particles by discarding distinctly poor-quality 2D classes to yield a partially cleaned particle stack of 303,048 particles. Ab-initio reconstruction of four volumes was performed to produce a single good 3D class and three poor 3D classes. Additional particle cleaning was performed using four parallel four-structure Heterogenous Refinement jobs, using the one good 3D class and two poor 3D classes derived from the previous Ab-initio Reconstruction job as inputs for the refinement. This parallel Heterogenous Refinement particle cleaning step was repeated three additional times, keeping the particles from the good classes for each subsequent cleaning steps, yielding a cleaned particle stack of 171,070 particles. The cleaned particles were aligned along their C2 symmetry axis using the Non-Uniform Refinement job ^47^ and then symmetry expanded to double the effective number of particles (342,140 particles). Here, some low-resolution speckles of EM density were visible on the periphery of the nucleosomal DNA. To determine if these speckles are Haspin bound to nucleosome or noise, two rounds of focused 3D classification were performed with a spherical focus mask centered on the area of speckled density to yield a discrete structure of Haspin bound to nucleosome. The structure was refined with Local Refinement to produce Haspin bound to nucleosome (30,638 particles 4.74 Å). This structure was used to project 2D templates for a template picking job in a subsequent processing workflow of a publication quality cryo-EM dataset of Hapsin bound to nucleosome. collected using a Titan Krios 300 kV electron microscope on a different grid produced in the same grid freezing session and under the same conditions as this test cryo-EM dataset.

The publication-quality dataset of Haspin bound to nucleosome collected on a Titan Krios 300 kV electron microscope on a grid produced in the same grid freezing session and under the same conditions as the test cryo-EM dataset. The publication-quality dataset was processed in cryoSPARC v4.2 ^46^. Exposures were imported with an EER upsampling factor of 2 and cropped to one-half their original resolution using Patch Motion Correction. The CTF correction was performed Patch CTF Correction. Poor quality micrographs were removed using Manually Curate Exposures, yielding 9,362 high-quality micrographs. A set of 2D templates from the test dataset of Haspin bound to nucleosome were used to perform particle picking using Template Picker and Inspect Picks, then extracted using Extract from Micrographs to yield an uncleaned particle stack (6,762,618 particles). One round of 2D Classification and Select 2D Classes was performed to remove junk particles by discarding distinctly poor-quality 2D classes to yield a partially cleaned particle stack (2,249,799 particles). Multiple parallel multi-structure Ab-initio reconstruction jobs were performed to produce one good 3D class and three poor 3D classes for subsequent heterogenous refinement particle cleaning. Particle cleaning was performed using four parallel four-structure Heterogenous Refinement jobs, using the one good 3D class and three poor 3D classes derived from the previous Ab-initio Reconstruction job as inputs for the refinement. This parallel Heterogenous Refinement particle cleaning step was repeated two additional times, keeping the particles from the good classes for each subsequent cleaning steps, yielding a cleaned particle stack of 900,411 particles. The cleaned particle stack was re-extracted and recentered using aligned shifts. Ab-initio Reconstruction and Homogenous Refinement jobs were performed to produce a structure showing some low-resolution speckles of EM density on the periphery of the nucleosomal DNA in two places on opposite sides of the nucleosome adjacent to the histone H3 tail. Focused 3D classification was performed with single mask file containing combined two spherical mask volumes each centered on one the two areas of speckled EM density. The output volumes showed structures of free nucleosome and structures of Haspin bound to nucleosome. The particles corresponding to Haspin bound to nucleosome were selected and refined with Homogenous Refinement then Local refinement. A second round of focused 3D classification was performed with a spherical mask centered on the Haspin density to produce structures of Haspin bound to nucleosome in two positions. Individual particle CTF was refined with local CTF refinement, image group CTF was refined with global CTF refinement, and the structures were refined with local refinement to produce Haspin bound to nucleosome in position 1 (152,199 particles, 3.01 Å), and Hapsin bound to nucleosome in position 2 (154,151 particles, 2.99 Å).

### Cryo-EM model building and refinement

Initial models of Haspin bound to nucleosome were constructed by using rigid body fitting of models for an unmodified nucleosome (PDB: 4ZUX)^48^, and Haspin (PDB: 4OUC) ^30^ into EM density maps using in ChimeraX ^49^, and then refined using all-atom flexible refinement with strong restraints in *Coot* 0.9.6 ^50^. All waters, ions, and small molecules were stripped from the original structures so that only the atoms corresponding to nucleosome and Haspin could be used for model building. Histone tails were extended where density was visible, and the nucleosomal DNA was extended from 147 bp to 157 bp to account for the extra-nucleosomal linker DNA present in this sample. The modeled Haspin includes residues 470-798. The models were further refined in PHENIX ^51^ using phenix.real_space_refine ^52^ and validated using the cryo-EM Comprehensive Validation module in PHENIX running MolProbity ^53^. Figures were generated with ChimeraX ^49^.

### Electrophoretic mobility shift assays

Binding reactions were prepared in 12 μL volumes by combining Haspin and nucleosome in binding buffer (20 mM HEPES pH 7.6, 150 mM NaCl, 5% glycerol, 1 mM DTT). Nucleosome was always added last. Prepared reactions were incubated on ice for 30 min to allow them to come to equilibrium. Prior to sample loading, 6% TBE gels were equilibrated by running at 150 V for 60 min at 4°C in 0.25X Tris-borate-EDTA (TBE) buffer. Then, 10 μL of each equilibrated binding reaction was loaded on the prepared gel and run at 150 V for 90 min at 4°C in 0.25X TBE buffer. Gels were stained in the dark on a rotating shaker for 20 min with SYBR Gold (Invitrogen) DNA-intercalating stain diluted to 1:5000, then imaged.

### Immunoblotting activity assays

Assays to test wild-type Haspin (465-798) and Haspin mutants (465-798) histone H3 threonine 3 (H3T3) phosphorylation activity were prepared in 20 μL volumes by combining Haspin and nucleosome in kinase buffer (20 mM HEPES pH 7.6, 150 mM KCl, 1 mM MgCl_2_, 1 mM DTT, 100 μM ATP, and 0.25 mg/mL bovine serum albumin (BSA) (GoldBio #A-420-250)). Nucleosomes was added last to initiate the reactions, which were incubated at room temp. Time points at 5, 15, and 45 minutes were collected by quenching 6 μL of each reaction in 1.5X lithium dodecyl sulfate (LDS) sample buffer. Prior to sample loading, Quenched reactions were separated on a 4-12% Bis-Tris Gel (Thermo Fisher) by SDS-PAGE then transferred onto polyvinylidene difluoride (PVDF) membrane using a Trans-Blot Turbo Transfer System (BioRad). Membranes were blocked with 5% milk in 1X Tris-buffered saline with 0.1% Tween-20 (TBST) buffer, probed with 1:5000 anti-H3T3ph (Sigma-Aldrich #04-746) or 1:5000 anti-H3 (Abcam #ab1791) primary antibodies in 5% milk in 1X TBST overnight at 4°C, then probed with 1:5000 anti-rabbit (Invitrogen #31460) horseradish peroxidase (HRP)- conjugated secondary antibody in 5% milk in 1X TBST for 1 hour at room temp. Membranes were stained with enhanced chemiluminescence (ECL) reagent and imaged.

### Cell lines and culture conditions

HEK293T cells were grown in Dulbecco’s Modified Eagle Medium (DMEM) (Corning Cellgro) containing 10% fetal bovine serum (Sigma), 100 U/mL penicillin, 100 U/mL streptomycin, and 2 mM L-glutamine and were maintained at 37 °C in a 5% CO2 atmosphere with 21% oxygen.

### Fluorescence microscopy

For fluorescence microscopy experiments, cells were grown to 75% confluency in a 6-well dish, then transfected with pcDNA5/FRT plasmids (Invitrogen, #V601020) expressing either enhanced green fluorescent protein (EGFP) alone, EGFP-Haspin (WT), or EGFP-Haspin (K569E, K759E, K761E, R772E) using Lipofectamine LTX transfection reagent (ThermoFisher, #15338500). Cells were transferred to 12-mm glass coverslips the day after and were fixed in MeOH at -20°C for 10 min on the third day. DNA was stained with 4′,6-diamidino-2-phenylindole (DAPI) in phosphate buffered saline (PBS) and cells were mounted with ProLong Gold Antifade reagent (Invitrogen).

Imaging was performed using a Zeiss Axio Observer 7 inverted microscope with Slidebook 2023 software (3i—Intelligent, Imaging Innovations, Inc.), CSU-W1 (Yokogawa) T1 50 μm Spinning Disk, and Prime 95B CMOS camera (Teledyne Photometrics) with a 63x plan-apochromat oil immersion objective with 1.4 NA. Mitotic cells were imaged with 0.25 μm z sections, deconvolved using the Slidebook Microvolution algorithm, and maximum intensity projected before analysis. Using FIJI software ^54^, mitotic chromosome regions were established from the DAPI channel, and median EGFP signal intensity was measured within the chromosome region (EGFP_DNA_) and for a 1 μm-wide cytoplasmic ribbon surrounding the DNA (EGFP_CYTO_). The ratio of intensity measurements was reported on a Log2 scale and plotted in GraphPad Prism.

### Foldseek protein structure homology search

The Foldseek^41^ Search Web Server (https://search.foldseek.com/search) was used with default parameters to search for structurally homologous proteins in the alphafold proteome. The C-terminal lobe of the Haspin kinase domain (residues 610-798) from the structure of Haspin bound to nucleosome in position 1 was used as a search query and the results were filtered to include only human proteins.

## DATA AVAILABILITY

Models and cryoEM maps were deposited in the Protein Data Bank (PDB) and Electron Microscopy Data Bank (EMDB) under the following accession codes: Haspin position 1 (PDB: 9B2S, EMDB: 44113), Haspin position 2 (PDB: 9B2T, EMDB: 44114), and Haspin local refinement (PDB: 9B2U, EMDB: 44115),

Raw cryoEM movies were deposited in the EMPIAR database under the accession code: EMPIAR-11971.

## ACKNOWLEDGEMENTS

We thank Duncan Sousa, Dazhong (David) Ding, and Kai Cai for their support with cryo-EM sample preparation and data collection at the Beckman Center for Cryo-EM at Johns Hopkins. We thank the Summer Academic Research Experience (SARE) Program at Johns Hopkins for providing us the opportunity to host Stacy Vasquez for a summer research experience in the Wolberger Lab. We thank the Wolberger lab for their insights and discussions on the manuscript.

This work was supported by the National Institute of General Medical Sciences (R35GM130393 to C.W., R01GM133897 to A.H., R01GM114119 to A.H.); and the National Cancer Institute (F31CA261154 to C.W.H., F31CA271743 to S.R., R01CA266199 to A.H.) of the National Institutes of Health.

---

**Figure S1.**
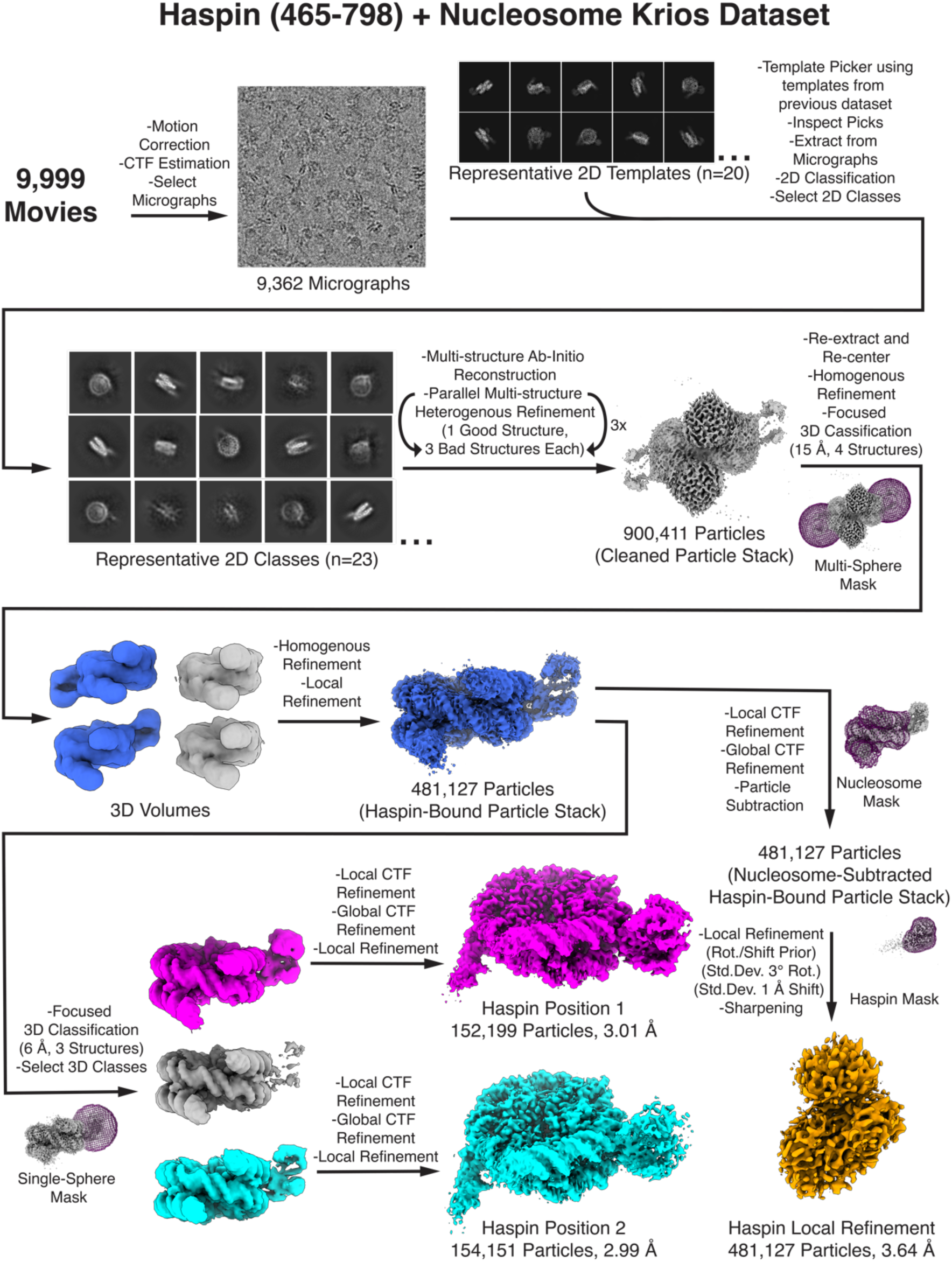
Cryo-EM data processing workflow for Haspin (465-798) bound to nucleosome.

**Figure S2.**
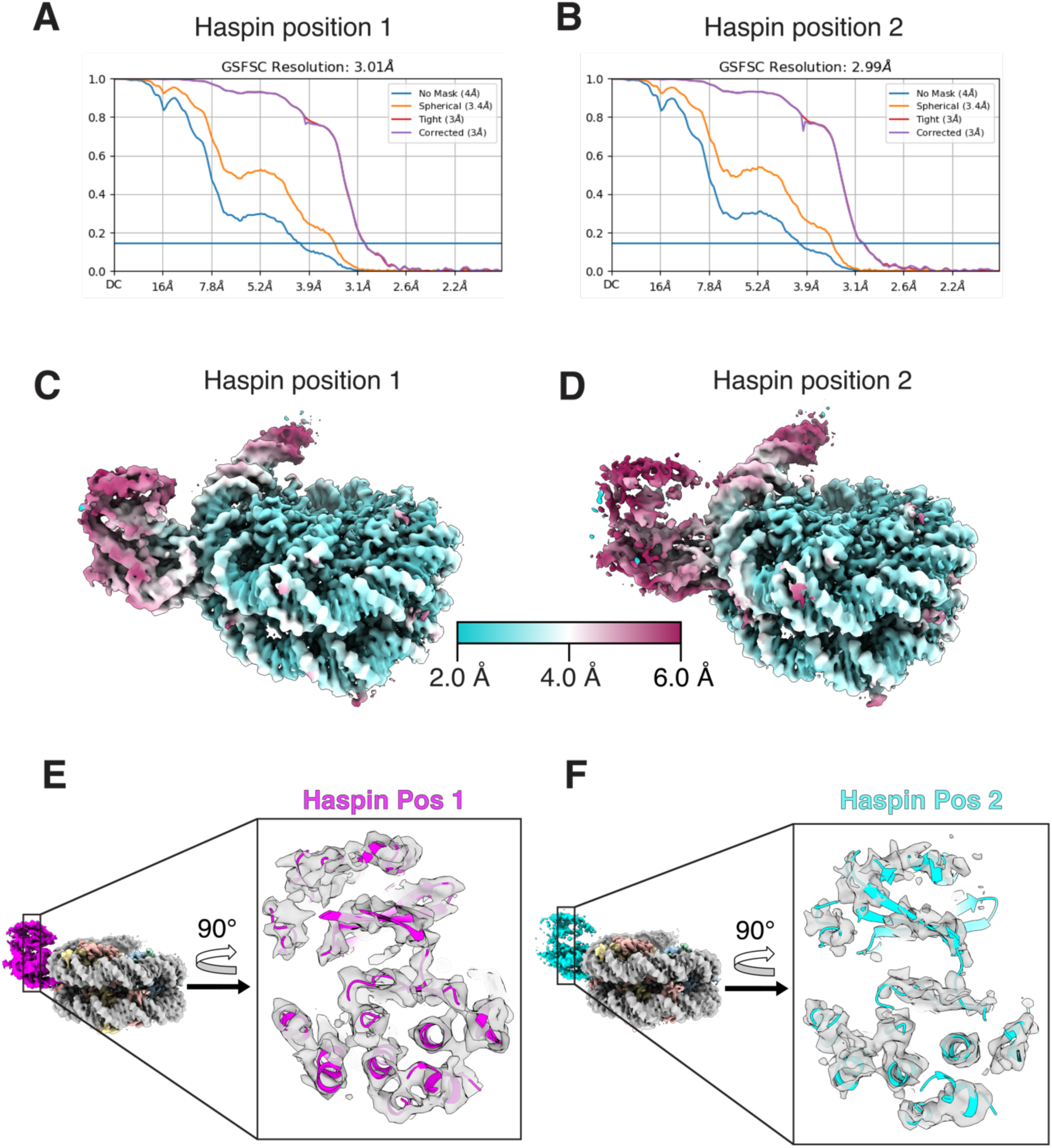
Global and local resolution evaluation of both cryo-EM maps. **A-B)** Fourier Shell Correlation (FSC) plots using fold-standard 0.143 cutoffs for both cryo-EM maps of Haspin bound to nucleosome. **C-D)** Local resolution estimation color depictions for both cryo-EM maps corresponding to Haspin bound to nucleosome. **E-F)** Cut-away slice view through the center of both Haspin cartoon models showing the fit to their respective cryo-EM maps.

**Figure S3.**
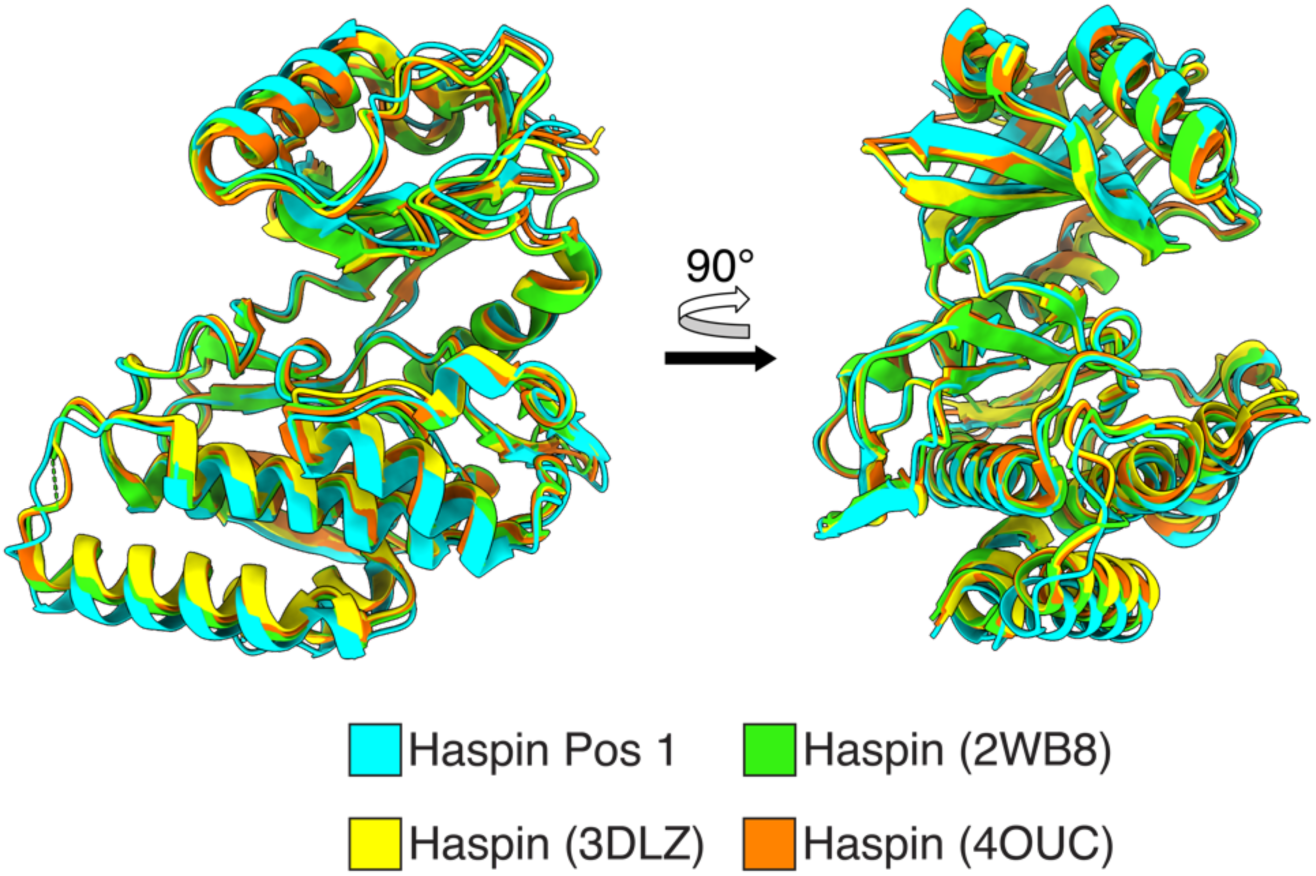
Haspin structural alignments. Structural alignments of cartoon models of Haspin (470-798) in position 1 over Haspin (468-798) (PDB: 2WB8) (Villa et al., 2009), Haspin (470-798) (PDB: 3DLZ) (Eswaran et al., 2009), and Haspin (470-798) (PDB: 4OUC) (Maiolica et al., 2014).

**Figure S4.**
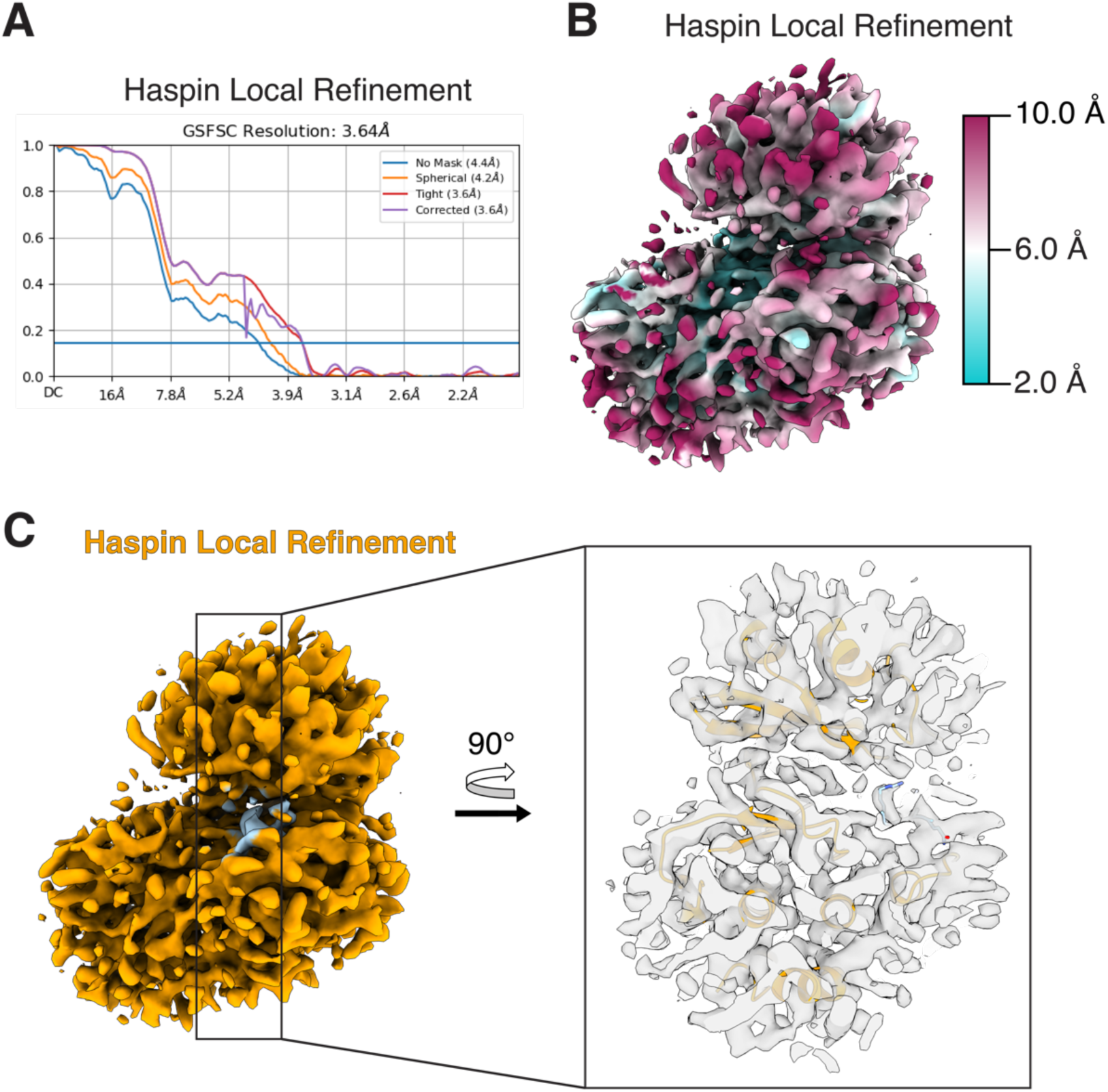
Global and local resolution evaluation of Haspin local refinement cryo-EM map. **A)** Fourier Shell Correlation (FSC) plot using fold-standard 0.143 cutoffs for the Haspin local refinement cryo-EM map of Haspin bound to nucleosome. **B)** Local resolution estimation color depictions for the Haspin local refinement cryo-EM map of Haspin bound to nucleosome. **C)** Cut-away slice view through the center of Haspin showing the fit of the model to the cryo-EM map.

**Figure S5.**
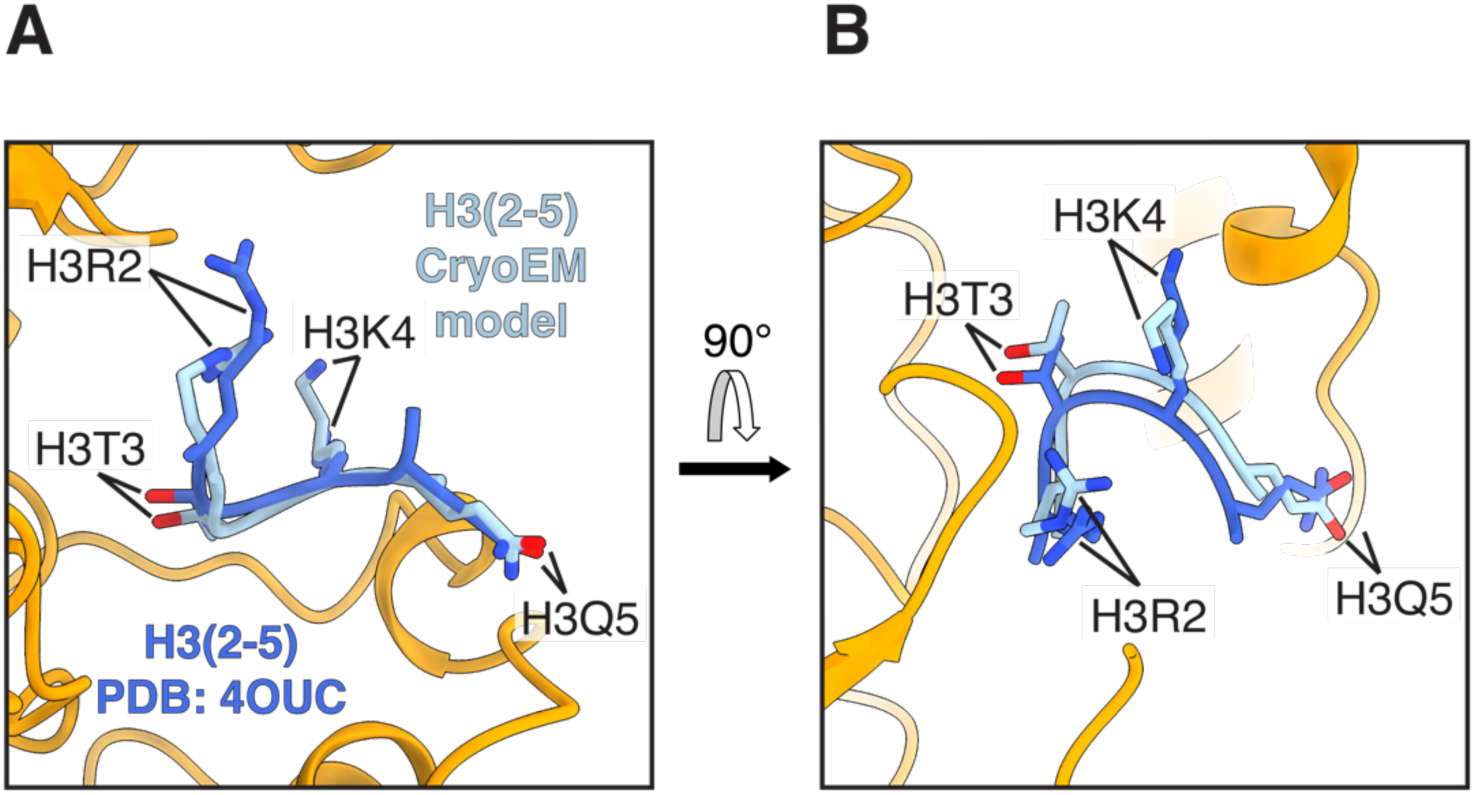
Cryo-EM structure of H3 tail bound to Haspin is similar to crystal structure of H3 peptide bound to Haspin. A previously reported crystal structure of Haspin (PDB: 4OUC) (Maiolica et al., 2014) with bound H3 peptide (royal blue) was superimposed over our cryo-EM map of locally-refined Haspin (orange) containing bound H3 tail (light blue). Structures are depicted in cartoon and atom representation and show the similarity in the binding position of H3.

## REFERENCES

1. Bannister, A.J. & Kouzarides, T. Regulation of chromatin by histone modifications. Cell research 21, 381–395 (2011).

2. Wang, F. & Higgins, J.M. Histone modifications and mitosis: countermarks, landmarks, and bookmarks. Trends in cell biology 23, 175–184 (2013).

3. Wilkins, B.J. et al. A cascade of histone modifications induces chromatin condensation in mitosis. Science 343, 77–80 (2014).

4. Yamagishi, Y., Honda, T., Tanno, Y. & Watanabe, Y. Two histone marks establish the inner centromere and chromosome bi-orientation. Science 330, 239–243 (2010).

5. Polioudaki, H. et al. Mitotic phosphorylation of histone H3 at threonine 3. FEBS letters 560, 39–44 (2004).

6. Qian, J., Lesage, B., Beullens, M., Van Eynde, A. & Bollen, M. PP1/Repo-man dephosphorylates mitotic histone H3 at T3 and regulates chromosomal aurora B targeting. Current Biology 21, 766–773 (2011).

7. Wang, F. et al. Histone H3 Thr-3 phosphorylation by Haspin positions Aurora B at centromeres in mitosis. Science 330, 231–235 (2010).

8. Kelly, A.E. et al. Survivin reads phosphorylated histone H3 threonine 3 to activate the mitotic kinase Aurora B. Science 330, 235–239 (2010).

9. Carmena, M., Wheelock, M., Funabiki, H. & Earnshaw, W.C. The chromosomal passenger complex (CPC): from easy rider to the godfather of mitosis. Nature reviews Molecular cell biology 13, 789–803 (2012).

10. Crosio, C. et al. Mitotic phosphorylation of histone H3: spatio-temporal regulation by mammalian Aurora kinases. Molecular and cellular biology 22, 874–885 (2002).

11. Welburn, J.P. et al. Aurora B phosphorylates spatially distinct targets to differentially regulate the kinetochore-microtubule interface. Molecular cell 38, 383–392 (2010).

12. Dai, J. & Higgins, J.M. Haspin: a mitotic histone kinase required for metaphase chromosome alignment. Cell cycle 4, 665–668 (2005).

13. Dai, J., Sultan, S., Taylor, S.S. & Higgins, J.M. The kinase haspin is required for mitotic histone H3 Thr 3 phosphorylation and normal metaphase chromosome alignment. Genes & development 19, 472–488 (2005).

14. Higgins, J.M. Haspin-like proteins: A new family of evolutionarily conserved putative eukaryotic protein kinases. Protein Science 10, 1677–1684 (2001).

15. Higgins, J. Structure, function and evolution of haspin and haspinrelated proteins, a distinctive group of eukaryotic protein kinases. Cellular and Molecular Life Sciences CMLS 60, 446–462 (2003).

16. Wang, F. et al. Haspin inhibitors reveal centromeric functions of Aurora B in chromosome segregation. Journal of Cell Biology 199, 251–268 (2012).

17. Dai, J., Sullivan, B.A. & Higgins, J.M. Regulation of mitotic chromosome cohesion by Haspin and Aurora B. Developmental cell 11, 741–750 (2006).

18. Markaki, Y., Christogianni, A., Politou, A.S. & Georgatos, S.D. Phosphorylation of histone H3 at Thr3 is part of a combinatorial pattern that marks and configures mitotic chromatin. Journal of cell science 122, 2809–2819 (2009).

19. Hadders, M.A. et al. Untangling the contribution of Haspin and Bub1 to Aurora B function during mitosis. Journal of Cell Biology 219, e201907087 (2020).

20. Kawashima, S.A., Yamagishi, Y., Honda, T., Ishiguro, K.-i. & Watanabe, Y. Phosphorylation of H2A by Bub1 prevents chromosomal instability through localizing shugoshin. Science 327, 172–177 (2010).

21. Liang, C. et al. Centromere-localized Aurora B kinase is required for the fidelity of chromosome segregation. Journal of Cell Biology 219(2020).

22. Kang, T.-H. et al. Mitotic histone H3 phosphorylation by vaccinia-related kinase 1 in mammalian cells. Molecular and cellular biology 27, 8533–8546 (2007).

23. Jeong, M.-W., Kang, T.-H., Kim, W., Choi, Y.H. & Kim, K.-T. Mitogen-activated protein kinase phosphatase 2 regulates histone H3 phosphorylation via interaction with vaccinia-related kinase 1. Molecular biology of the cell 24, 373–384 (2013).

24. Cartwright, T.N. et al. Dissecting the roles of Haspin and VRK1 in histone H3 phosphorylation during mitosis. Scientific Reports 12, 11210 (2022).

25. Tanaka, H. et al. Identification and characterization of a haploid germ cell-specific nuclear ProteinKinase (haspin) in spermatid nuclei and its effects on somatic cells. Journal of Biological Chemistry 274, 17049–17057 (1999).

26. Tanaka, H. et al. Cloning and characterization of human haspin gene encoding haploid germ cell-specific nuclear protein kinase. Molecular human reproduction 7, 211–218 (2001).

27. Villa, F. et al. Crystal structure of the catalytic domain of Haspin, an atypical kinase implicated in chromatin organization. Proceedings of the National Academy of Sciences 106, 20204–20209 (2009).

28. Eswaran, J. et al. Structure and functional characterization of the atypical human kinase haspin. Proceedings of the National Academy of Sciences 106, 20198–20203 (2009).

29. Hanks, S.K. & Hunter, T. The eukaryotic protein kinase superfamily: kinase (catalytic) domain structure and classification 1. The FASEB journal 9, 576–596 (1995).

30. Maiolica, A. et al. Modulation of the chromatin phosphoproteome by the Haspin protein kinase. Molecular & cellular proteomics 13, 1724–1740 (2014).

31. Barski, A. et al. High-resolution profiling of histone methylations in the human genome. Cell 129, 823–837 (2007).

32. Quadri, R., Sertic, S. & Muzi-Falconi, M. Roles and regulation of Haspin kinase and its impact on carcinogenesis. Cellular signalling 93, 110303 (2022).

33. De Antoni, A., Maffini, S., Knapp, S., Musacchio, A. & Santaguida, S. A small-molecule inhibitor of Haspin alters the kinetochore functions of Aurora B. Journal of Cell Biology 199, 269–284 (2012).

34. Wotring, L.L. & Townsend, L.B. Study of the cytotoxicity and metabolism of 4-amino-3-carboxamido-1-(β-d-ribofuranosyl) pyrazolo [3, 4-d] pyrimidine using inhibitors of adenosine kinase and adenosine deaminase. Cancer research 39, 3018–3023 (1979).

35. Fedorov, O. et al. A systematic interaction map of validated kinase inhibitors with Ser/Thr kinases. Proceedings of the National Academy of Sciences 104, 20523–20528 (2007).

36. Amoussou, N.G., Bigot, A., Roussakis, C. & Robert, J.-M.H. Haspin: a promising target for the design of inhibitors as potent anticancer drugs. Drug Discovery Today 23, 409–415 (2018).

37. Horn, V. & van Ingen, H. Recognition of nucleosomes by chromatin factors: Lessons from data-driven docking-based structures of nucleosome-protein complexes, (IntechOpen, 2018).

38. McGinty, R.K. & Tan, S. Recognition of the nucleosome by chromatin factors and enzymes. Current Opinion in Structural Biology 37, 54–61 (2016).

39. Kale, S., Goncearenco, A., Markov, Y., Landsman, D. & Panchenko, A.R. Molecular recognition of nucleosomes by binding partners. Current opinion in structural biology 56, 164–170 (2019).

40. Edayathumangalam, R.S., Weyermann, P., Gottesfeld, J.M., Dervan, P.B. & Luger, K. Molecular recognition of the nucleosomal “supergroove”. Proceedings of the National Academy of Sciences 101, 6864–6869 (2004).

41. van Kempen, M. et al. Fast and accurate protein structure search with Foldseek. Nat Biotechnol 42, 243–246 (2024).

42. Luger, K., Rechsteiner, T.J. & Richmond, T.J. Preparation of nucleosome core particle from recombinant histones. in Methods in enzymology, Vol. 304 3–19 (Elsevier, 1999).

43. Lowary, P. & Widom, J. New DNA sequence rules for high affinity binding to histone octamer and sequence-directed nucleosome positioning. Journal of molecular biology 276, 19–42 (1998).

44. Makde, R.D., England, J.R., Yennawar, H.P. & Tan, S. Structure of RCC1 chromatin factor bound to the nucleosome core particle. Nature 467, 562–566 (2010).

45. Dyer, P.N. et al. Reconstitution of nucleosome core particles from recombinant histones and DNA. in Methods in enzymology, Vol. 375 23–44 (Elsevier, 2003).

46. Punjani, A., Rubinstein, J.L., Fleet, D.J. & Brubaker, M.A. cryoSPARC: algorithms for rapid unsupervised cryo-EM structure determination. Nature methods 14, 290–296 (2017).

47. Punjani, A., Zhang, H. & Fleet, D.J. Non-uniform refinement: adaptive regularization improves single-particle cryo-EM reconstruction. Nature methods 17, 1214–1221 (2020).

48. Morgan, M.T. et al. Structural basis for histone H2B deubiquitination by the SAGA DUB module. Science 351, 725–728 (2016).

49. Pettersen, E.F. et al. UCSF ChimeraX: Structure visualization for researchers, educators, and developers. Protein Science 30, 70–82 (2021).

50. Emsley, P., Lohkamp, B., Scott, W.G. & Cowtan, K. Features and development of Coot. Acta Crystallographica Section D: Biological Crystallography 66, 486–501 (2010).

51. Liebschner, D. et al. Macromolecular structure determination using X-rays, neutrons and electrons: recent developments in Phenix. Acta Crystallographica Section D: Structural Biology 75, 861–877 (2019).

52. Afonine, P.V. et al. Real-space refinement in PHENIX for cryo-EM and crystallography. Acta Crystallographica Section D: Structural Biology 74, 531–544 (2018).

53. Williams, C.J. et al. MolProbity: More and better reference data for improved all-atom structure validation. Protein Science 27, 293–315 (2018).

54. Schindelin, J., et al. Fiji: an open-source platform for biological-image analysis. Nature methods 9, 676–682 (2012).

